# The potential of 4-Methylumbelliferone to be repurposed for treating liver fibrosis

**DOI:** 10.1101/2025.05.19.654778

**Authors:** Xi Chen, Huiqiao Li, Yanru Deng, Jieyi Meng, Shangang Zhao, Clavia Ruth Wooton-Kee, Xia Gao, Bingning Dong, Dongyin Guan, Chaodong Wu, Philipp E. Scherer, Yi Zhu

**Author notes:** Please address correspondence to Yi Zhu.

## Abstract

4-Methylumbelliferone (4-MU) is the active component of hymecromone, a choleretic and antispasmodic drug with an excellent safety profile. In rodent studies, high doses of 4-MU are also used to inhibit the production of hyaluronan (HA), a biomarker of liver fibrosis. Further, 4-MU shows excellent efficacy in inhibiting liver fibrosis of different etiologies in animal studies, eliciting interest in its repurposing for this condition. However, 4-MU’s mechanism of action, and whether it inhibits liver fibrosis by impeding HA synthesis, remains unclear. Using several transgenic mouse models with HA overproduction or degradation in different types of liver cells, we found that both directions of perturbation reduced liver fibrosis levels. In addition, degrading HA via hyaluronidase PH20 overexpression impaired liver function manifested by increased serum aminotransferase (ALT) activity levels. These findings challenge both the role of HA modulation in 4-MU’s action and the strategy of targeting HA to treat liver fibrosis. Additional mouse models also excluded the possibility that 4-MU modulates intestinal farnesoid X receptor (FXR) to inhibit liver fibrosis. Ablation of gut microbiota partially abolishes 4-MU’s anti-liver fibrosis effect. However, the anti-liver fibrosis effect of 4-MU was lost in the lower-dose group, and the high dose’s effect in reducing ALT disappeared over time despite its liver fibrosis-reducing effect. Based on these findings, we argue that the lack of efficacy of 4-MU at a translatable dose and lack of a clear mechanism that allows further improvement of 4-MU’s efficacy make 4-MU impractical for development as an anti-liver fibrosis treatment.

**Highlights:**

- 4-MU inhibits MASH fibrosis.
- Both HA overproduction and digestion in the liver reduce liver fibrosis.
- Intestinal epithelial cell FXR is not required for 4-MU to reduce liver fibrosis.
- The gut microbiota is partially responsible for 4-MU’s antifibrotic effect in the liver.
- 4-MU lacks the potential to be further developed into anti-liver fibrosis treatment.

## Introduction

4-MU (7-hydroxy-4-methylcoumarin) is a coumarin derivative, with positions seven hydroxylated and four methylated. It is the active component of hymecromone, a choleretic and antispasmodic drug approved in Europe and Asia. In healthy human subjects, hymecromone increases common bile duct diameter after a standard meal [1] and delays postprandial contraction of the common bile duct [2], which forms the foundation for its use as a choleretic and antispasmodic agent.

4-MU is also widely used as an inhibitor of hyaluronan (HA) synthesis. HA is a non-sulfated glycosaminoglycan (GAG) polymer composed of repeating disaccharide units of D-glucuronic acid (GlcUA) and N-acetyl-D-glucosamine (GlcNAc). It is a major component of the extracellular matrix (ECM), with high abundance in connective, epithelial, and dermal tissues [3, 4]. 4-MU inhibits HA synthesis by conjugating with GlcUA to deplete the cellular UDP-GlcUA pool required for HA synthesis [5] as well as via an unknown mechanism to downregulate *HAS2*expression [6]. Notably, there is no evidence that 4-MU directly inhibits the enzymatic activity of any HAS isoforms.

Nonalcoholic steatohepatitis (NASH) is a subtype of nonalcoholic fatty liver disease (NAFLD) characterized by inflammation and liver fibrosis [7–9]. Revently, NASH has been renamed metabolic dysfunction-associated steatohepatitis (MASH) to emphasize the contribution of metabolic dysfunction to the pathogenesis of the disease [10]. MASH is currently the most common form of chronic liver disease worldwide [11, 12], and liver fibrosis in MASH patients remains the strongest predictor of adverse clinical outcomes [13]. The U.S. Food and Drug Administration (FDA) just approved the first drug, Madrigal’s resmetirom, an oral liver-targeted thyroid hormone receptor-beta (THRβ) agonist, for treating NASH (the term used by the FDA to describe this condition) [14]. However, the effect size is moderate-resmetirom improve liver fibrosis by at least one stage, with no worsening of the NAFLD activity score in 25% of the treatment group compared to 15% of the placebo group [14]. Thus, it is necessary to identify more effective treatment for NASH, especially for liver fibrosis.

HA, together with procollagen III N-terminal peptide (PIIINP) and tissue inhibitor of metalloproteinase 1 (TIMP1), is used to calculate the enhanced liver fibrosis (ELF) score, a non-invasive diagnostic tool for detectingsevere fibrosis in patients with chronic hepatitis B, hepatitis C, and HIV. HA has also been used as a surrogate marker for fibrosis in NASH/MASH clinical trials [15]. Beyond serving as a biomarker, HA is suspected to play a direct role in liver fibrogenesis by promoting ECM secretion from the hepatic stellate cells (HSCs) [16]. This connection had led to an interest in testing whether 4-MU could prevent or slow liver fibrogenesis. In animal experiments, 4-MU has shown excellent efficacy in improving liver fibrosis caused by CCl_4_, bile duct ligation (BDL), or choline-deficient diet treatment [16–21]. However, whether oral 4-MU ameliorates liver fibrosis by suppressing hepatic HA synthesis remains unclear. Using several transgenic mouse models with increased HA synthesis or degradation, we showed that both HA overproduction and digestion protected mice from liver fibrogenesis, highlighting the complex role of HA in liver fibrogenesis and raising the possibility that 4-MU may act through alternative mechanisms to inhibit liver fibrogenesis.

To explore this possibility, we investigated other mechanisms through which 4-MU might exert its antifibrotic effects. However, findings from additional animal models raised reasonable doubts about the involvement of a single specific pathway mediating the antifibrotic effects of high-dose 4-MU. This lack of a clear mechanism poses a significant barrier to developing more potent 4-MU-based derivatives for the treatment of liver fibrosis.

## Results

### Only extremely high-dose 4-MU reduces liver fibrosis

In our previous study, 5% 4-MU supplemented with a high-fat diet (HFD) significantly improved glucose tolerance in C56BL/6J mice [22]. Liver samples from these mice were used to quantify the expression of the liver fibrosis-related genes. After a 5-week HFD treatment, the expression of the major collagen genes *Col1a1* and *Col1a2* was suppressed by 4-MU supplementation, while *Acta2*, which encodes smooth muscle α-actin—a key component of stress fibers within cells—did not change (Fig. 1A). As it is well known that HFD alone is not sufficient to induce liver fibrosis, we treated a new cohort of mice with CDAHFD, with or without 0.2% 4-MU or 5% 4-MU. After four weeks of treatment, 5% 4-MU significantly suppressed *Col1a1*, *Col3a1*, and *Acta2* gene expression, whereas 0.2% 4-MU failed to suppress these markers (Fig. 1B). When another cohort of mice were treated for 8 weeks with 5% 4-MU CDAHFD, a similar level of suppression of the liver fibrosis genes was observed (Supplementary Fig. 1A). The liver HA content was reduced by 72.9% (Fig. 1C). Picrosirius red (PSR) staining showed significant suppression of liver fibrosis by 4-MU (Fig. 1D). Liver collagen content also decreased by 46.7% (Fig. 1E). Only 5% 4-MU treatment significantly reduced ALT and AST levels after 4 weeks of treatment (Fig. 1F), but the reduction in ALT and AST levels was lost after 8 weeks of treatment (Supplementary Fig. 1B).

**Fig. 1.**
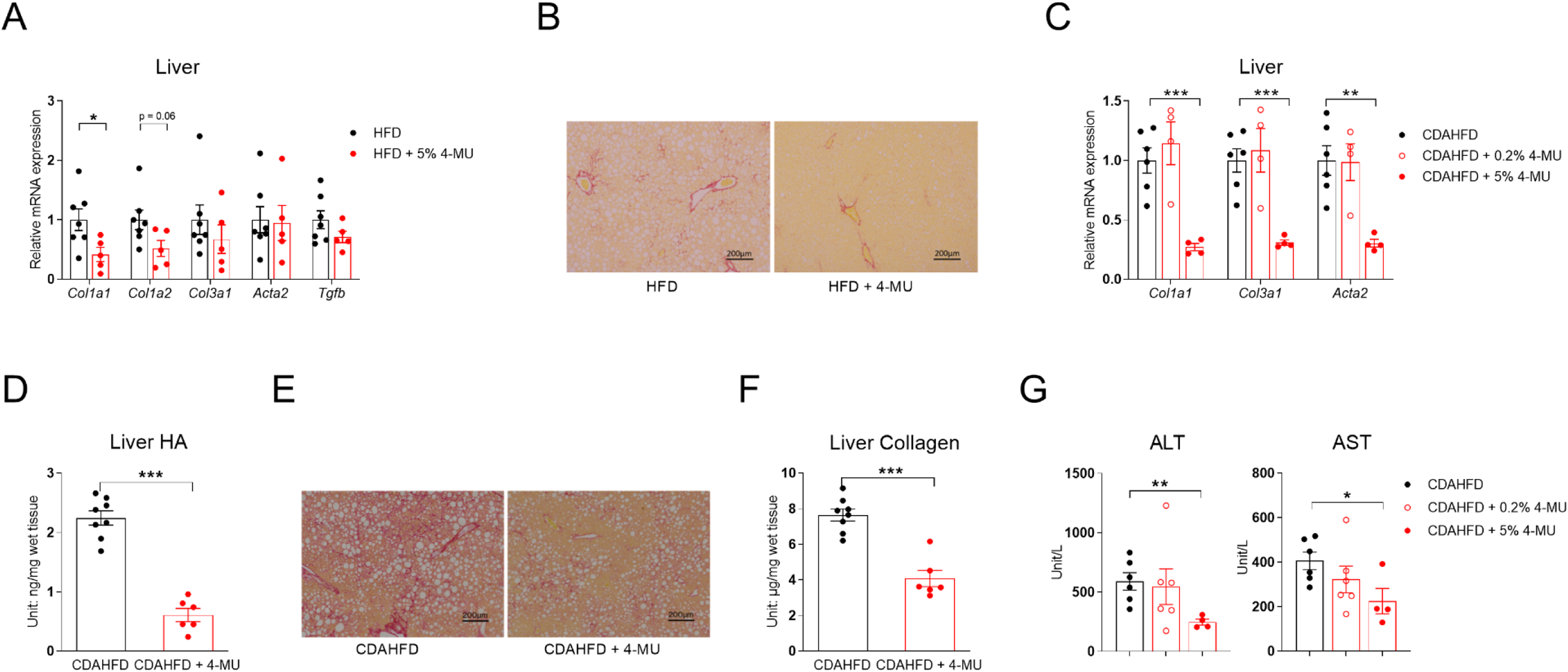
4-MU reduces liver fibrosis in CDAHFD-treated C57BL/6J mice. **(A)** Relative mRNA expression of fibrosis-related genes in the livers of C57BL/6J mice kept on 5-week HFD or HFD with 5% 4-MU, *n* = 5–7. **(B)** Relative mRNA expression of fibrosis-related genes in the livers of C57BL/6J mice kept on CDAHFD or CDAHFD with 0.2% or 5% 4-MU for 4 weeks, *n* = 4–6. **(C)** Liver hyaluronic acid (HA) content of C57BL/6J mice kept on CDAHFD or CDAHFD with 5% 4-MU for 8 weeks. **(D)** The representative liver PSR-staining images from the mice used in panel D, scale bar = 500 μm. **(E)** Quantification of liver collagen content in the mice from panels D and D; for panels D–F, *n* = 6–8. **(F)** Serum ALT and AST levels of C57BL/6J mice kept on CDAHFD or CDAHFD with 0.2% or 5% 4-MU for 4 weeks, *n* = 4–6. All data are presented as mean ± SEM; * *p* < 0.05, ** *p* < 0.01, *** *p* < 0.001.

### Intrahepatic production of HA reduces liver fibrosis

4-MU is commonly used to inhibit HA synthesis [5, 6], and we observed a robust reduction in hepatic HA contents in mice treated with 4-MU (Fig. 1C). To evaluate the causality of HA in liver fibrosis, we first tested whether forced expression of *Has2*, which would increase HA production, aggravates liver fibrosis. We crossed Lrat-rtTA [23] and TRE-*Has2* [22] mice to generate compound transgenic mice Lrat-rtTA::TRE-*Has2* (referred to as stellate cell *Has2*, or SHS mice), which overexpress *Has2* specifically in HSCs after treating with doxycycline (Fig. 2A). Unexpectedly, SHS mice displayed reduced *Col1a1* and *Col1a2* expression (Fig. 2B), slightly lower PSR-staining intensity (Fig. 2C) and significantly decreased liver collagen content on a liver fibrosis-inducing diet (Fig. 2D). Notably, H&E staining showed no overall difference in morphology (Supplementary Fig. 2), and ALT and AST levels did not differ (Fig. 2E). The immortalized human hepatic stellate cell line LX-2 transfected with *Has2* expressed lower levels of *COL1A1*, indicating either a cell-autonomous effect of mouse *Has2* expression or an autocrine effect of the produced HA on HSC collagen gene expression and liver fibrogenesis (Fig. 2F). To distinguish between these two possibilities, we used liver tissue from Alb-cre/Rosa-rtTA/TRE-*Has2* mice (referred to as Liv-*Has2* TG mice), as in our previous study [22], and found reducedexpression of *Col1a1* and *Col1a2* (Fig. 2G) without changes in ALT and AST levels [22]. These data demonstrated that extracellular HA, regardless of whether it is produced by HSC or adjacent hepatocytes, exerts a similar effect in suppressing liver fibrogenesis, supporting the idea that ECM HA plays an anti-fibrogenic role in the liver.

**Fig. 2.**
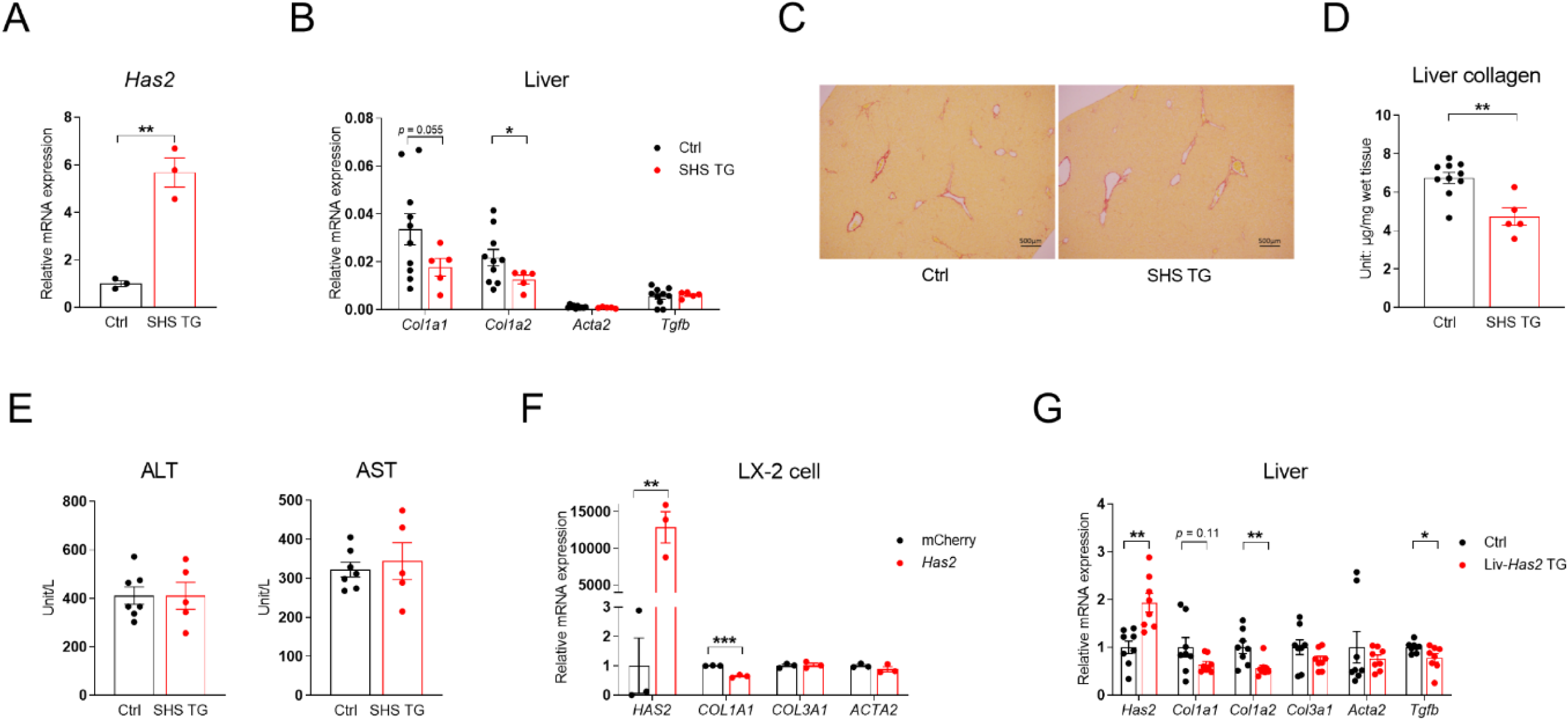
Intrahepatic production of HA reduces liver fibrosis. **(A)** Relative mRNA expression of *Has2* in isolated HSCs from control (Ctrl) and SHS TG mice, *n* = 3. **(B)** Relative mRNA expression of hepatic fibrosis-related genes in the livers of Ctrl and SHS TG mice treated with a 4-week dox200 MCD diet. **(C)** Representative liver PSR-staining images from mice in panel B, scale bar = 500 μm. **(D)** Liver collagen content of Ctrl and SHS TG mice from panel B; for panels B–D, *n* = 5–10. **(E)** Serum ALT and AST levels of Ctrl and SHS TG mice fed with 4-week dox200 MCD diet, *n* = 5–7. **(F)** Relative mRNA expression of *Has2* and fibrosis-related genes in LX-2 cells overexpressing *mCherry* as Ctrl or *Has2, n* = 3. **(G)** Relative mRNA expression of *Has2* and fibrosis-related genes in the Ctrl and Liv-*Has2* TG mice livers, *n* = 8. All data are presented as mean ± SEM; * *p* < 0.05, ** *p* < 0.01, *** *p* < 0.001.

### Systemic PEG-PH20 treatment or whole-body PH20 overexpression reduces liver fibrosis

To complement the *Has2* overexpression study, degradation of HA was enhanced using hyaluronidase PH20. PH20, encoded by *Spam1*, is a sperm-specific, secreted, glycosylphosphatidylinositol-anchored cell surface hyaluronidase that functions at neutral pH [24]. It demonstrated the highest activity in reducing culture medium HA concentration when transfected and expressed in HEK293 cells among the four hyaluronidases tested (Fig. 3A). Using bovine testis-derived PH20, we showed that it dose-dependently reduces HA content in LX-2 cell culture medium independent of TGF-β treatment (Fig. 3B). As expected, TGF-β significantly increased the expression of *COL1A1*, *ACTA2,* and *TGFB*, indicative of fibrotic activatio. The highest dose of PH20 reduced the expression of *COL1A1*, while all doses reduced *ACTA2* expression in LX-2 cells treated with TGF-β (Fig. 3C).

**Fig. 3.**
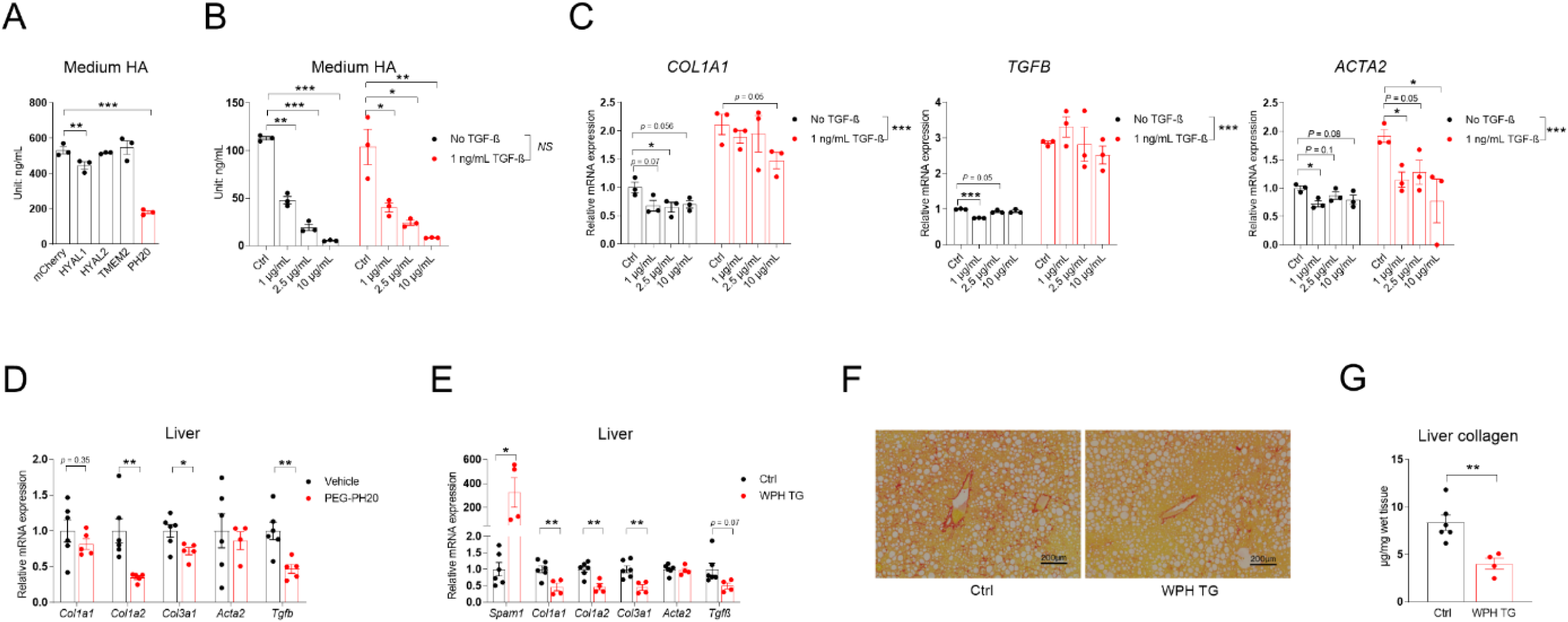
Systemic PEG-PH20 treatment or whole-body PH20 overexpression reduces liver fibrosis. **(A)** Culture medium HA content of HEK293 cells transfected with *mCherry* or different mouse hyaluronidase genes, *n* = 3. **(B)** Medium HA content of LX-2 cells treated with different concentrations of bovine testis PH20 in the presence or absence of 1 ng/mL TGF-β, *n* = 3. **(C)** Relative mRNA expression of COL1A1, *TGFB,* and *ACTA2* in LX-2 cells from panel B. **(D)** Relative mRNA expression of fibrosis-related genes in the livers of diet-induced obese mice injected with vehicle or 10 µg/g body weight of PEG-PH20, *n* = 5–6. **(E)** Relative mRNA expression of *Spam1* and fibrosis-related genes in the livers of control or WPH TG mice fed with CDAHFD for 8 weeks. **(F)** Representative liver PSR-staining images of mice in panel E, scale bar = 500 μm. **(G)** Liver collagen content of control and WPH TG mice from panel E; for panels E–G, *n* = 4–6. All data are presented as mean ± SEM; * *p* < 0.05, ** *p* < 0.01, *** *p* < 0.001.

In mice, pegylated human PH20 (PEG-PH20) treatment improved skeletal muscle blood perfusion, insulin sensitivity, and overall glucose metabolism in diet-induced obese (DIO) mice [25]. Gene expression analysis of the livers from these mice showed suppression of *Col1a2*, *Col3a1,* and *Tgfb* expression following PEG-PH20 treatment (Fig. 3D). As PEG-PH20 was no longer available for replication in a CDAHFD-induced liver fibrosis model, we generated TRE-PH20 mice to allow for doxycycline-inducible, cell-specific PH20 overexpression. A founder with doxycycline inducibility and no off-target expressionwas selected (Supplementary Fig. 3A). The functionality of *Spam1* overexpression in reducingHA levels was confirmed in adipose tissue, which has a higher basal HA concentration than liver in healthy mice (Supplementary Fig. 3B). The selected TRE-PH20 founder was crossed with whole-body R26-M2rtTA mice (Jax, 006965) to generate double-transgenic mice (referred to as WPH). WPH mice showed a more than 300-fold increase in *Spam1* expression in the liver after treatment with chow diet containing 200 mg/kg doxycycline (Fig. 3E). When treated with CDAHFD (containing 200 mg/kg doxycycline) for 8 weeks, these mice showed a drastic decrease in *Col1a1*, *Col1a2*, *Col3a1*, and *Tgfb* expression, whereas *Acta2* expression remained unchanged (Fig. 3E). PSR staining (Fig. 3F) and live collagen content quantification (Fig. 3G) suggested a robust reduction in fibrosis in WPH livers.

### Stellate cell and hepatocyte PH20 overexpression reduces liver fibrosis

To further determine PH20’s effect on liver fibrosis, TRE-PH20 mice were crossed with Lrat-rtTA mice [23] to generate Lrat-rtTA::TRE-PH20 mice, which overexpress PH20 specifically in HSC, and this mouse line was named stellate cell PH20 (SPH) (Fig. 4A). When treated with CDAHFD containing doxycycline, SPH miceshowed significant decreases in hepatic *Col1a1*, *Acta2*, and *Tgfb* expression (Fig. 4B), along with a 33.4% reduction in liver collagen content (Fig. 4C). These data show that the digestion of HA surrounding HSCs reduces liver fibrosis.

**Fig. 4.**
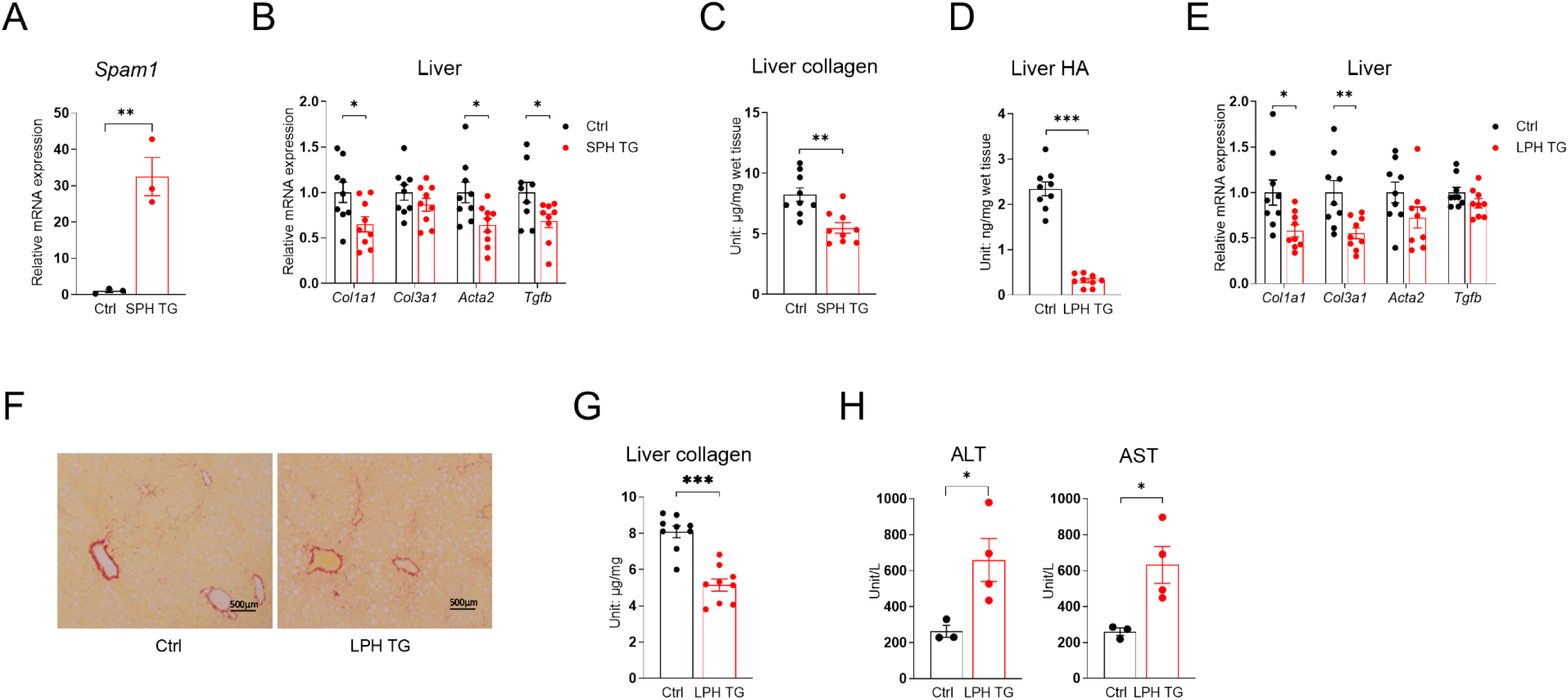
Hepatic stellate cell and hepatocyte PH20 overexpression reduces liver fibrosis, and 4-MU further reduces liver fibrosis in hepatocyte PH20-overexpressing mice. **(A)** Relative mRNA expression of *Spam1* in HSCs of control (Ctrl) and SPH TG mice, *n* = 3. **(B)** Relative mRNA expression of fibrosis-related genes in the livers of Ctrl and SPH TG mice fed with dox200 CDAHFD for 8 weeks. **(C)** Liver collagen content of Ctrl and SPH TG mice from panel B; for panels B–C, *n* = 9. **(D)** Liver HA content of Ctrl and LPH TG mice kept on dox200 CDAHFD for 8 weeks. **(E)** Relative mRNA expression of fibrosis-related genes in the livers of Ctrl and LPH TG mice. **(F)** The representative liver PSR-staining images of mice in panel D, scale bar = 500 μm. **(G)** Liver collagen content of Ctrl and LPH TG mice from panel D; for panels D–G, *n* = 8–9. **(H)** Serum ALT and AST levels of Ctrl and LPH TG mice treated with dox200 CDAHFD for 8 weeks, *n* = 3–4. All data are presented as mean ± SEM; * *p* < 0.05, ** *p* < 0.01, *** *p* < 0.001.

Since PH20 is secreted and functions in the extracellular space [24], PH20 secreted by hepatocytes adjacent to HSCs is expected to exert a similar or even more significant effect on ECM HA degradation and suppression of HSC activation and liver fibrogenesis, given the predominant number and mass of hepatocytes in the liver. As expected, in Alb-Cre::Rosa-rtTA::TRE-PH20 (referred to as liver PH20 transgenic mice, abbreviated as LPH mice) (Supplementary Fig. 3A and 4), PH20 robustly digested liver HA (Fig. 4D). Consistent with this, we observed a similar reduction in liver fibrosis gene expression (Fig. 4E), as well as significantly diminished PSR staining (Fig. 4F) and liver collagen content (Fig. 4G). Notably, the expression of PH20 in hepatocytes drastically increased serum ALT and AST levels (Fig. 4H), cautioning against the use of PH20 to treat liver fibrosis.

### 4-MU does not ameliorate liver fibrosis through activating intestinal FXR

Given the complex role of HA in liver fibrogenesis, we sought to uncover other mechanisms through which 4-MU may reduce liver fibrosis. 4-MU is a substrate inhibitor of UGT, competing with UDP for conjugation to glucuronic acid (GA) and thereby reducing the formation of UDP-GA, which is required for UGT-catalyzed conjugation and excretion of BAs from IECs [26]. IEC-retained intracellular BAs activate FXR to induce FGF15 expression and secretion, which binds to hepatocyte FGFR4 to suppress bile acid metabolism, hepatic glucose metabolism, and lipogenesis [27–29]. FGF15, a mouse ortholog of human FGF19, may ameliorate liver fibrosis by improving hepatic and systemic metabolism [30]. Furthermore, a novel intestinal FXR agonist improves liver fibrosis through modulation of the gut-liver axis [31]. Therefore, another possibility for how 4-MU could regulate liver fibrogenesis involves increasing FGF15 levels by inhibiting UGT-catalyzed glucuronidation and excretion of Bas from IECs.

4-MU treatment in CDAHFD-fed mice significantly increased serum FGF15 levels compared to CDAHFD control mice (Fig. 5A), which correlated with the suppression of several cytochrome P450 genes responsible for BAs’ synthesis (Fig. 5B) and with a reduction in serum BAs (Fig. 5C). To assess the requirement of intestinal FXR for the antifibrotic effects of 4-MU, we crossed Vil-CRE-ERT2 mice with *Fxr* flox mice to generate Vil-CRE-ERT2::*Fxr*^fl/fl^ mice (referred to as VFX-KO mice). A single tamoxifen injection (100 mg/kg, delivered intraperitoneally) robustly induced *Fxr* deletion in IECs (Supplementary Fig. 5). To overcome potential IEC regeneration, we treated control and VFX-KO mice with tamoxifen weekly during the CDAHFD feeding. Tamoxifen treatment completely abolished IEC *Fxr* expression in VFX-KO mice, whereas 4-MU treatment did not affect *Fxr* expression (Fig. 5D). Notably, intestinal *Fxr* deletion did not affect *Col1a1* expression, while 4-MU suppressed *Col1a1* expression, regardless of the presence or absence of *Fxr* (Fig. 5D and 5E). Similarly, PSR staining showed that IEC-specific deletion of *Fxr* did not affect fibrosis, while 4-MU suppressed fibrosis in both control and VFX-KO mice (Fig. 5F). Quantification of liver collagen levels showed a similar pattern (Fig. 5G). All of these data suggest that 4-MU does not ameliorate liver fibrosis by activating intestinal FXR.

**Fig. 5.**
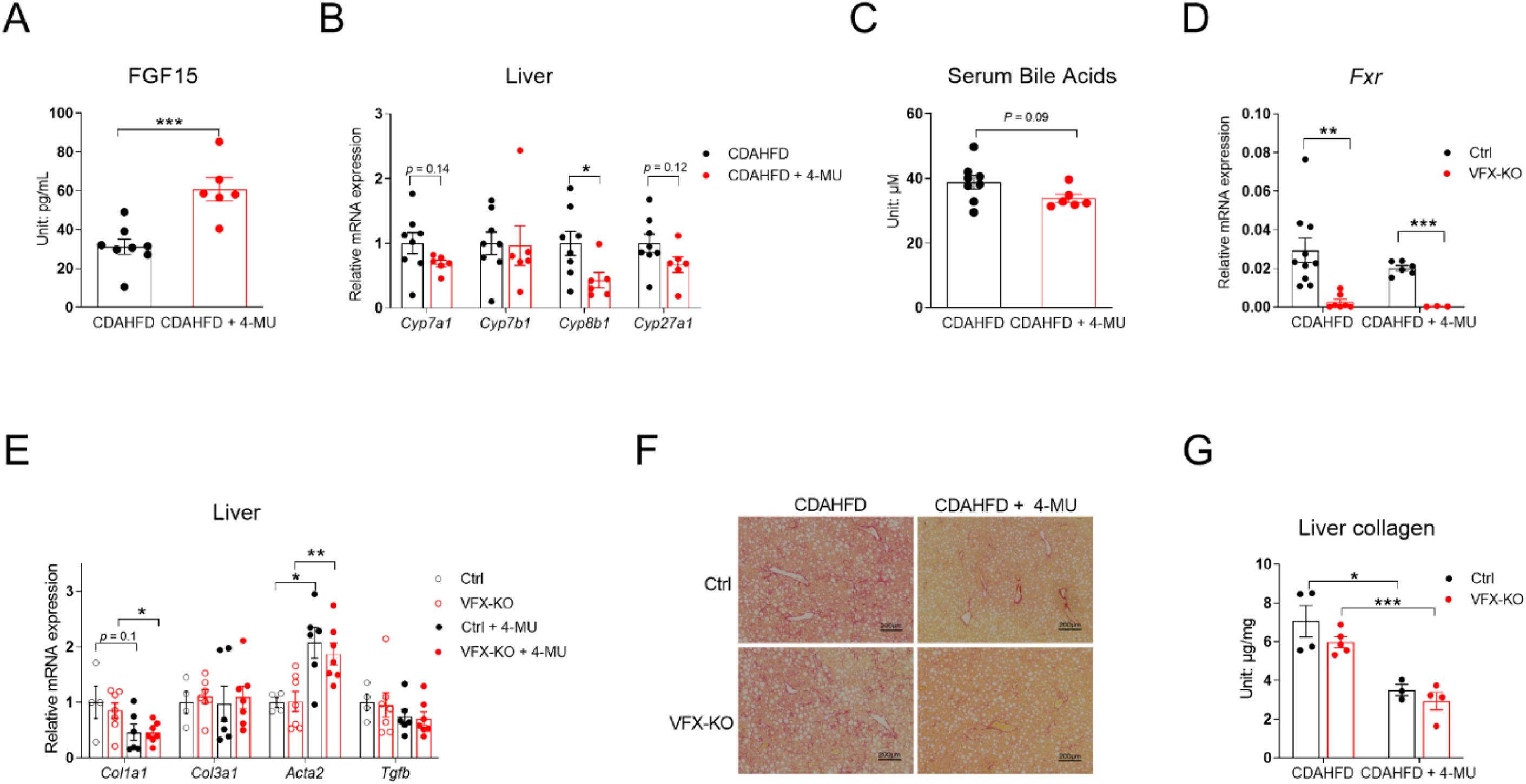
4-MU does not ameliorate liver fibrosis by activating intestinal FXR. **(A)** Serum FGF15 levels in mice treated with 8-week CDAHFD with or without 5% 4-MU. **(B)** Relative mRNA expression of bile acid metabolism-related genes in the livers of mice from panel A. **(C)** Serum bile acid levels of mice from panel A; for panels A–C, *n* = 6–8. **(D)** Relative mRNA expression of *Fxr* in control (Ctrl) and VFX-KO mice maintained on CDAHFD with or without 5% 4-MU for 4 weeks, *n* = 3–10. (**E)** Relative mRNA expression of fibrosis-related genes in the livers of Ctrl and VFX-KO mice fed CDAHFD with or without 5% 4-MU for 4 weeks. **(F)** Representative liver PSR-staining images of mice in panel E. **(G)** Liver collagen content of Ctrl and VFX-KO mice, *n* = 3–5. All data are presented as mean ± SEM; * *p* < 0.05, ** *p* < 0.01, *** *p* < 0.001.

### Hepatic *Fxr* deletion significantly reduced liver fibrosis, and 4-MU further reduced the fibrosis

Next, we tested whether 4-MU’s efficacy is dependent on hepatic FXR. Albumin-Cre mice were crossed with *Fxr* flox mice to generate Alb-Cre::*Fxr*^fl/fl^ (LFX-KO) mice. Following CDAHFD treatment, LFX-KO mice showed markedly enlarged gallbladders and yellowish mucosa (Fig. 6A). Gallbladder enlargement was also observed in LFX-KO mice treated with HFD, but to a lesser extent, and their livers also appeared less steatotic (Supplementary Fig. 6). Deletion of *Fxr* significantly elevated liver and serum BAs content (Fig. 6B), as previously reported [32]. However, ileum BAs levels were reduced in LFX-KO mice fed CDAHFD (Fig. 6B), likely due to reduced hepatic *Abcb11* expression (encoding bile salt export pump, known as BSEP) (Fig. 6C). 4-MU treatment generally had no effect on liver or ileum BAs levels, except that it significantly reduced serum BAs levels of control mice (Fig. 6B). In addition to *Abcb11*, deletion of *Fxr* in the liver also led to near-complete suppression of *Cyp7b1*, which encoded 25-hydroxycholesterol 7-alpha-hydroxylase in the BA synthesis pathway (Fig. 6C). LFX-KO mice showed significantly reduced *Col1a1* expression, and 4-MU treatment further reduced its expression (Fig. 6D). PSR staining and liver collagen quantification exhibited a similar pattern: hepatic *Fxr* deletion reduced liver fibrosis, but 4-MU treatment further reduced it (Fig. 6E, 6F). However, the deletion of *Fxr* in the liver significantly elevated ALT and AST levels, while 4-MU treatment did not affect them (Fig. 6G). Altogether, these data suggest that 4-MU does not require hepatic FXR to alleviate liver fibrogenesis and that deletion of hepatic FXR leads to liver damage.

**Fig. 6.**
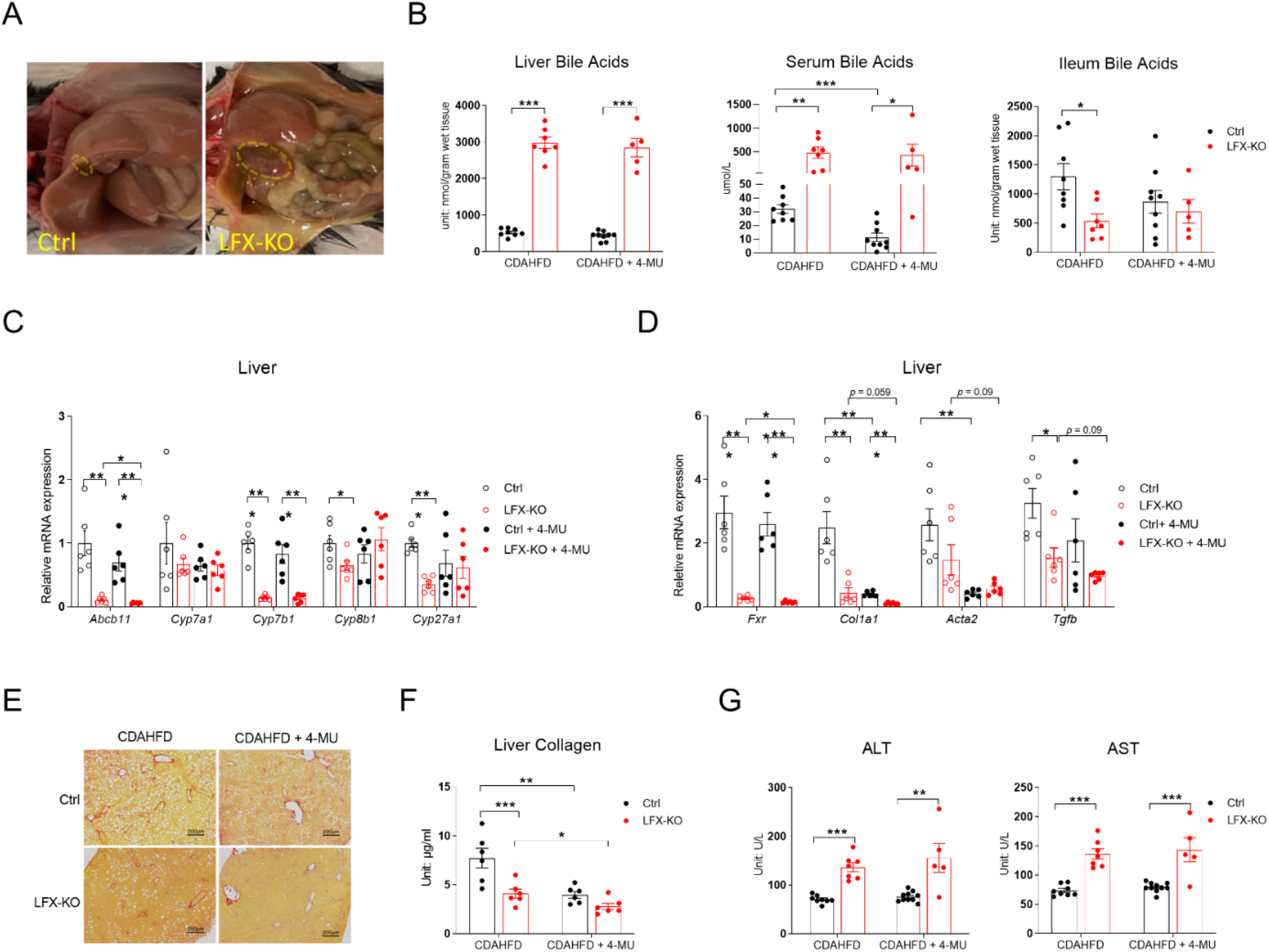
Hepatic *Fxr* deletion significantly reduces liver fibrosis, and 4-MU further reduces liver fibrosis. **(A)** Gross appearance of livers from control (Ctrl) and LFX-KO mice treated with 8-week CDAHFD. **(B)** Liver, serum, and ileum BAs levels of Ctrl and LFX-KO mice treated with 8-week CDAHFD with or without 5% 4-MU. **(C)** Relative mRNA expression of *Abcb11* and BAs-synthesis genes in the livers of Ctrl and LFX-KO mice treated with 8-week CDAHFD with or without 5% 4-MU. **(D)** Relative mRNA expression of *Fxr* and fibrosis-related genes in the livers of Ctrl and LFX-KO mice from panel C. **(E)** Representative PSR staining of mice in panel C. **(F)** Liver collagen content of Ctrl and LFX-KO mice from panel C. **(G)** Serum ALT and AST levels of WT and LFX-KO mice from panel B. Two cohorts of mice were used: for panels B and G, *n* = 5– 10; for panels C–F, *n* = 6. All data are presented as mean ± SEM; * *p* < 0.05, ** *p* < 0.01, *** *p* < 0.001.

### Microbiota ablation by antibiotic treatment partially blocks 4-MU’s antifibrotic effects

Importantly, 4-MU inhibits not only mammalian UGTs but also bacterial UGTs [33]. Given the critical role of the gut microbiota in liver fibrogenesis [34, 35], we sought to test whether 4-MU modulates liver fibrosis development by reshapeing the microbiota. We used a combination of four antibiotics [36], which completely eradicated the gut bacteria within three weeks (Supplementary Fig. 7A). A new cohort of mice were then treated with antibiotics concurrently with CDAHFD. The mice did not develop diarrhea, and the intestinal length was not affected by CDAHFD or antibiotic water treatment (Supplementary Fig. 7B). Ablation of the gut microbiota alone did not affect *Col1a1*, *Col3a1,* and *Tgfb* expression, but it partially reversed 4-MU’s suppression of those genes (Fig. 7A). PRS staining of liver sections showed a more pronounced reversal of 4-MU-mediated liver fibrosis improvement upon gut microbiota ablation (Fig. 7B). Quantification of liver collagen levels revealed that antibiotic treatment alone did not affect liver fibrosis but partially blocked 4-MU’s suppression of liver fibrogenesis (Fig. 7C), a pattern similar to *Col1a1* expression. Although quantification of bacterial species by 16S sequencing showed that 4-MU treatment did not affect the species abundance or diversity (Supplementary Fig. 7C–F), it did significantly reduced the abundance of proteobacteria (Fig. 7D and E), a phylum known to release lipopolysaccharide (LPS) and increase gut inflammation and liver fibrogenesis [37–40].

**Fig. 7.**
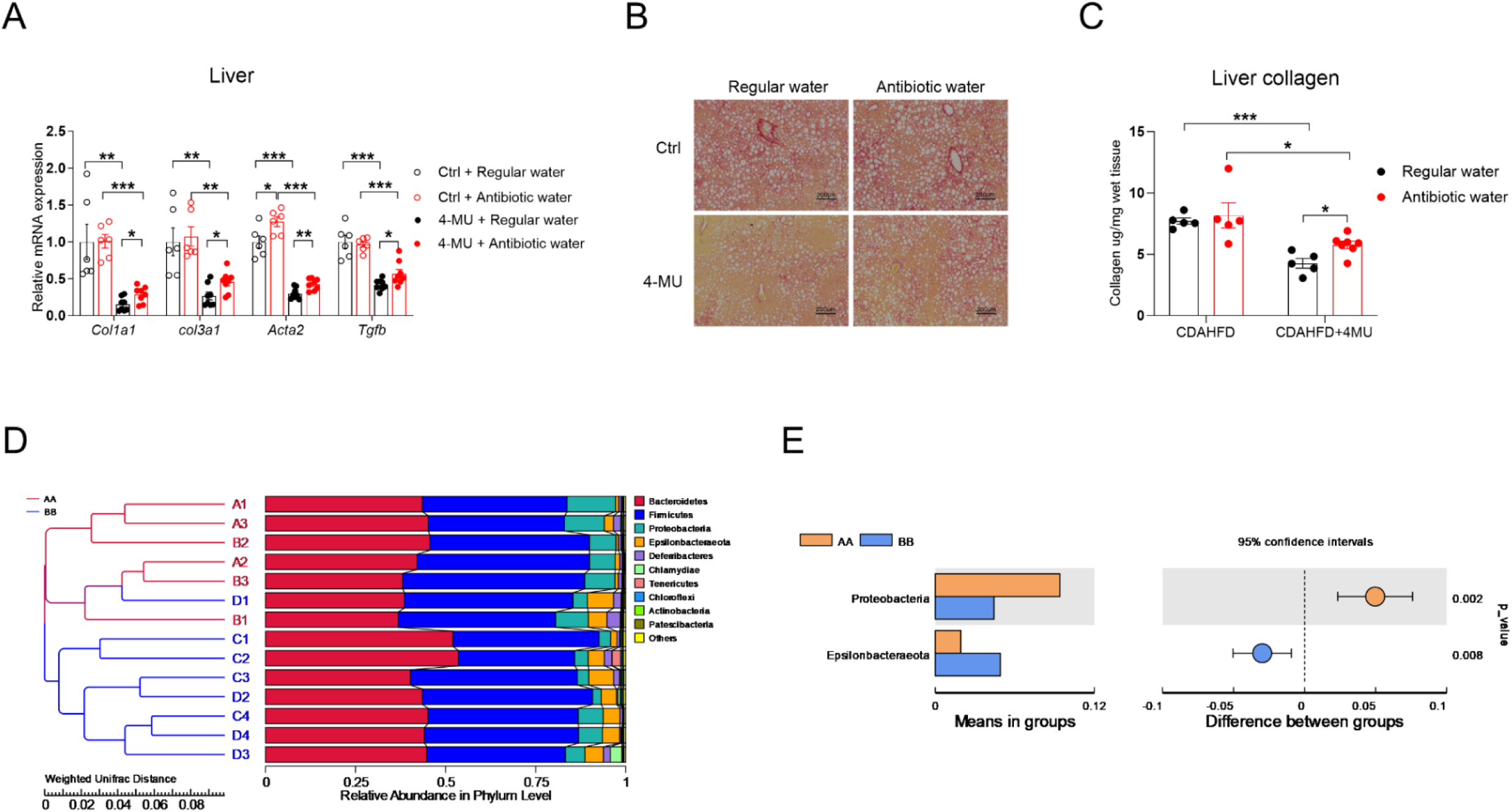
Antibiotic treatment partially blocks 4-MU’s antifibrotic effect. **(A)** Relative mRNA expression of fibrosis-related genes in WT mice treated with 8-week CDAHFD with or without 5% 4-MU, which were provided with regular water or antibiotic water, *n* = 6–8. **(B)** Representative liver PSR staining from panel A, scale bar = 500 μm. **(C)** Liver collagen content of WT mice with various treatments, *n* = 5–7. **(D)** Quantification of bacterial species by 16S sequencing; AA/red line: control group, comprising A1–A3 and B1–B3, two separate cohorts given the same treatment but harvested at different times; BB/blue line: 4-MU group, comprising C1– C4 and D1–D4, two separate cohorts given the same treatment but harvested at different times. Accession to cite for these SRA data: PRJNA1130997. **(E)** The relative abundance of proteobacteria and Epsilonbacteraeota. AA: vehicle, BB: 4-MU treatment. All data are presented as mean ± SEM; * *p* < 0.05, ** *p* < 0.01, *** *p* < 0.001.

## Discussion and Conclusions

The pathogenesis of NASH/MASH involves whole-body metabolic dysregulation, along with a complex signaling disequilibrium and functional impairment of multiple types of cells in the liver [41, 42]. These led to a lack of consensus on druggable targets for the disease. In addition, patients with NASH/MASH show great interpatient variability in the expression of selected drug targets [43], further contributing to a historically high failure rate in clinical trials. Examples of high-profile failures include the FXR agonist obeticholic acid, the dual PPARα/δ agonist elafibranor, the CCR2/CCR5 inhibitor cenicriviroc, and the ASK1 inhibitor selonsertib, representing the challenges in safely targeting master liver nuclear receptors, inflammation, and cell death to treat NASH/MASH [44, 45].

The nomenclature change of NASH to MASH highlights the contribution of metabolic dysregulation to the development of this liver disease [10]. Serendipitously, the first FDA-approved NASH treatment, resmetirom, is a thyroid hormone receptor-beta agonist, which functions by regulating lipid metabolism and reducing liver fat accumulation [46]. However, it shows mediocre efficacy in reducing liver fibrosis, and thus, better therapeutic options are still needed for patients suffering from this devastating disease. 4-MU has attracted considerable attention because it effectively reduces liver fibrosis in animal models and is approved in many countries with an excellent safety profile.

However, the biggest caveat in those animal studies is that the mice had to receive extremely large doses of 4-MU. Most studies, including the high-dose group in our study, put mice on a 5% 4-MU-containing diet, which translates to 5,000 mg/kg body weight per day, assuming mice weighing 30 grams consum 3 grams of food per day. Using the body surface area normalization method for dose translation from animals to humans [47], this dose equals 405.4 mg/kg in human patients, which means a 70 kg patient needs to take 28.4 grams of 4-MU. The use of this extremely high 4-MU dose was rooted in the belief that 4-MU inhibits HA synthesis by conjugating to glucuronic acid to prevent the formation of UDP-GlcUA and thus deplete the cellular UDP-GlcUA pool for HA synthesis [5]. UDP and UDP-GlcUA are abundantly present in hepatocytes [48], thus, a very high dose of 4-MU is required to compete with UDP to form 4-MU-GlcUA. However, even with this high dose, whether it efficiently inhibits HA synthesis and reduces HA levels in all HA-producing tissues, like adipose tissues, remains questionable [22]. There has been one study examining 4-MU’s effect on HA levels in humans. Three different oral doses (1.2 grams, 2.4 grams, and 3.6 grams) were tested, and only the 1.2-gram cohort showed a statistically significant reduction in serum HA levels from the baseline, while no change occurred in the 2.4-gram and 3.6-gram cohorts [49], raising concerns about the reproducibility of the findings in the 1.2-gram cohort.

While there is no doubt that HA levels significantly increase in fibrotic livers and in serum samples from patients with severe fibrosis, whether HA contributes to pathogenesis remains controversial. In a healthy liver, HA is only detected on sinusoids at low levels and is absent on the endothelial surface of post-sinusoidal venules or in the liver parenchyma [50]. HSC, the cell type responsible for collagen secretion and liver fibrogenesis, is the primary cell type contributing to HA synthesis, especially after being activated and transformed into myofibroblasts [16]. Thus, it is possible that an increase in HA levels in fibrotic livers is just a reflection of the activation of HSCs and the active proliferation of transformed HSCs.

To dissect HA’s role in liver fibrogenesis, we used doxycycline-inducible liver cell type-specific *Has2* and *Spam1*-overexpressing animals, which promote HA synthesis in *Has2* overexpressed cells or HA degradation around *Spam1* overexpressed cells, respectively. Our data demonstrate that both HA synthesis and degradation reduce liver fibrogenesis. In particular, Lrat-*Has2* transgenic animals showed slightly reduced liver fibrosis, in contrast to the enhanced liver fibrosis levels observed in Acta2-*Has2* mice [16]. This likely stems from the different promoters used in the studies: *Lrat* is active in quiescent HSCs, whereas *Acta2* is only active in activated HSCs and myofibroblasts. It is possible that HA synthesis protects quiescent cells from activation, which is supported by the in vitro culture of the human HSC cell line LX-2 cells, showing that overexpression of mouse *Has2* increases HA production and protects LX-2 cells from TGFβ-induced increases in *COL1A1* expression. However, after HSC activation, HA may support myofibroblast proliferation, contributing to upregulated liver fibrosis levels in Acta2-*Has2* mice. The pro-proliferative roles of HA are well supported by the literature on many other cell types, especially stem cells or stem-like cells [51, 52]. Working on the same pathway, digestion of HA by PH20 is expected to inhibit the proliferation of activated HSC, resulting in lower levels of liver fibrosis, as seen in our several *Spam1* overexpressing mouse models. Of note, hepatocyte *Spam1* overexpression increased AST and ALT levels, aruguing against the use of PH20 enzyme (encoded by *Spam1*) to digest HA as an antifibrotic treatment.

Aside from testing whether 4-MU inhibits liver fibrosis via inhibiting HA synthesis, we also tested whether 4-MU modulates bile acid metabolism and FGF15/19 secretion to inhibit liver fibrogenesis. The rationale was that the gastrointestinal system extracts approximately 40% of 4-MU [53], and that UGT in intestinal epithelial cells (IECs) plays an important role in BAs’ glucuronidation and excretion. Given this context, it is possible that 4-MU inhibits UGTs to prevent BAs’ glucuronidation and excretion. Retained cellular BAs then activate FXR and promote FGF15/19 expression and secretion to subsequently protects the liver from fibrogenesis. However, inducible deletion of *Fxr* in IEC did not affect liver fibrogenesis, and 4-MU still suppressed liver fibrogenesis in IEC-specific *Fxr* knockout mice.

4-MU is also a bacterial UGT inhibitor [33] and can be fermented by some bacteria [54], potentially alternating gut microbiota in mice. Using an antibiotic cocktail, we efficiently ablated the gut microbiota and showed that this partially abolished 4-MU’s anti-liver fibrotic effects, echoing the vital role of gut microbiota in the maintenance of liver health [34, 55]. 16S sequencing of feces revealed a significant reduction of proteobacteria species—Gram-negative bacteria with the presence of LPS in their outer membrane—after 4-MU treatment. This change in proteobacteria is expected to reduce gut inflammation to alleviate liver fibrogenesis [37–40]. However, since low-dose 4-MU failed to alleviate liver fibrosis, whether the approved dose can modulate gut microbiota in patients remains in doubt.

Finally, despite a consistent reduction in liver fibrosis with high-dose 4-MU treatment, improvements in liver function—as assessed by AST/ALT levels or *Acta2* expression—have been inconsistent. For example, after 8 weeks of 4-MU treatment in combination with CDAHFD, there was no difference in AST and ALT levels, suggesting no functional benefit of 4-MU despite a significant reduction in liver fibrosis. Across different experiments, we consistently observed reduced *Col1a1* expression and decreased tissue collagen content, consistent with the fact that fibrosis primarily consists of secreted extracellular collagen I fibers. In contrast, α-smooth muscle actin reflects cellular stress, and *Acta2* expression has been variable across animal models. For instance, an increase in *Acta2* expression was observed in both control and VFX-KO mice after 4-MU treatment, which may reflect increased hepatocyte stress due to the combined effects of tamoxifen injection and impaired liver detoxification caused by 4-MU. Specifically, 4-MU-mediated depletion of UDP-GlcUA in hepatocytes may interfere with detoxification and the elimination of waste compounds [48]. Ultimately, it remains possible that the reduction in *Col1a1* expression seen with high-dose 4-MU may simply reflect impaired HSC function with reduced collagen synthesis capacity.

In summary, only a very high dose of 4-MU exerts an antifibrotic effect on the mouse liver. The mechanism is likely multi-pronged, including improvement of metabolism [22] and modulation of gut microbiota. However, there is no clear route to improving 4-MU’s efficacy in those aspects due to the lack of a precise mechanism by which 4-MU improves them.

## Materials and Methods

### Mice

Our study mainly examined male mice because male animals exhibited less variability in phenotype. For the examination of long-term treatment in VFX mice, we used female mice because male mice were intolerant of tamoxifen treatment at the dose we used. TRE-*Has2* [22] and TRE-PH20 (also TRE-*Spam1*) were generated by subcloning mouse cDNA into the pTRE vector (Clontech) with a rabbit β-globin 3’-UTR. Linearized DNA was injected into an oocyte, and founders were identified through PCR. The founders were then crossed with a tissue-specific rtTA mouse to select a founder with tissue-specific transgene induction after doxycycline treatment and without leaking transgene expression in the nontarget tissue, or without doxycycline treatment. Other mice were obtained from the Jaxson laboratory or published before: Adipoq-rtTA mice (Jax, 033448) [56], Albumin-*Cre* transgenic (Jax, 003574) [57], Rosa26-loxP-STOP-loxP-rtTA (Jax, 005572) [58], Lrat-rtTA [23], *Fxr* floxed (Jax, 028393) [32], and Vil1-Cre/ERT2 (Jax, 020282) [59]. Tamoxifen (Sigma-Aldrich, T5648-1G) was dissolved in corn oil and injected into mice intraperitoneally to induce *Fxr* deletion. All animals were kept under a 12-hour light-dark cycle in a temperature-controlled environment. Mice were given free access to water and fed one of the following: CDA-HFD containing 5% 4-MU, CDA-HDF containing 200 mg/kg doxycycline, and CDA-HDF containing 200 mg/kg doxycycline + 5% 4-MU, which were custom-made based on the L-Amino Acid Diet with 45 kcal% Fat with 0.1% Methionine, No Added Choline and 1% Cholesterol diet (Research Diets, A16092003). To ablate the gut microbiota, mice were treated with broad-spectrum antibiotics in drinking water for four weeks (ampicillin, 1 g/L; neomycin sulfate, 1 g/L; metronidazole, 1 g/L; vancomycin, 500 mg/L) [36]. All treatments were done in male mice unless otherwise specified, starting around 10 weeks of age. Mice were genotyped in house using SYBR-based qPCR or at Transnetyx using TaqMan-based qPCR.

### Cell culture

LX-2 cells, an immortalized human HSC cell line [60], were purchased from Sigma-Aldrich (SCC064). LX-2 cells were cultured in high-glucose DMEM medium (Thermo Fisher, 10569010) containing 2% fetal bovine serum (Gibco, 10082147) and 1% penicillin-streptomycin (Sigma-Aldrich, P4333, 100 ml). LX-2 cells were transfected with the CMV-rtTA plus TRE-*mCherry* or TRE-*Has2* plasmids. 1μg/ml doxycycline was mixed with fresh complete medium and added to the transfected cells one day after transfection to induce gene expression. The cells were harvested 24 h after doxycycline addition for RT-qPCR analysis. For the TGF-β stimulation experiment, 1 ng/mL human TGF-β (Sigma-Aldrich, SRP3171) was used to activate LX-2 cells, and bovine testis hyaluronidase (Sigma-Aldrich, H3506) was used at the doses indicated in the text. After 24 h, the medium was collected for HA measurement and cells were harvested for RT-qPCR analysis.

HEK-293 cells, a human embryonic kidney cell line, were purchased from the American Type Culture Collection (ATCC; CRL-1573). HEK-293 cells were cultured in high-glucose DMEM containing 10% fetal bovine serum and 1% penicillin-streptomycin. Cells were transfected with CMV-rtTA plus TRE-*mCherry*, TRE-*Hyal1,* TRE-*Hyal2*, TRE-*Tmem2*, or *TRE-Spam1* plasmids. The following day, 1μg/ml doxycycline was mixed with fresh complete medium and added to the transfected cells to induce gene expression. The medium was collected for HA measurements after 24 h of incubation.

### Histology

After euthanasia, the tissues were immediately excised from mice, fixed overnight in 10% PBS-buffered formalin, and stored in 50% ethanol. Tissues were sectioned (5 µm), rehydrated, and stained with hematoxylin and eosin (H&E) or picrosirius red (PSR) at the Pathology Core at BCM. Microscopic images were obtained with a ZEISS Axioscan scanner.

### HA extraction and quantification

The procedure was performed as previously described [22], and the extracted HA was quantified using an ELISA kit (R&D Systems, DHYAL0), with a sample digested with bovine testis hyaluronidase overnight as the negative control. The tissue HA content was normalized to the wet weight used for extraction.

### qRT-PCR

RNA was isolated from frozen tissues by homogenization in TRIzol reagent (Invitrogen, 15596018), as previously described [61]. RNA concentration was quantified using a NanoDrop spectrophotometer. Using a reverse transcription kit (Bio-Rad), 1 µg of RNA was used to transcribe the cDNA. qRT-PCR primers were obtained from the Harvard PrimerBank [62] and are listed in Supplementary Table 3. Quantitative real-time PCR (qRT-PCR) was carried out using 2X Universal SYBR Green Fast qPCR Mix (ABclonal, RK21203). The mRNA levels were calculated using the comparative threshold cycle method and normalized to the gene *Rps16*.

### Bile acid extraction and quantification

Approximately 20 mg of liver, intestine, feces samples, or a whole gallbladder were homogenized in 0.2 ml 75% ethanol for 30 seconds. The homogenate was incubated in a 50°C water bath for 2 hours, with vortexing every 15–30 minutes. Then, the samples were spun for 10 min at 6,000 × g and the supernatant was collected. Extracted samples and serum were further diluted, if necessary, and assayed using the Mouse Total Bile Acids Assay Kit (Crystal Chem, 80471).

### Hepatic collagen extraction and quantification

Collagen quantification was conducted using a Total Collagen Assay Kit (Abcam, ab222942) following the manufacturer’s protocol. In brief, the collagen I standard was hydrolyzed via alkaline hydrolysis and subsequently diluted to different concentrations of the standard, as specified. Approximately 20 mg of fresh liver tissue was harvested and hydrolyzed. The samples were diluted with ddH_2_O to ensure their concentrations were within the standard curve range. Equal volumes of the sample hydrolysate and standard were added to a 96-well plate and evaporated by heating at 65°C to obtain a crystalline residue. The oxidation reagent mix, developer, and DMAB concentrate were added sequentially, with each addition followed by an incubation period at the corresponding temperature. Absorbance was measured at 560 nm using a microplate reader, and the final concentration was calculated by multiplying the absorbance by the dilution factor.

### FGF15 ELISA

Serum FGF15 levels were measured using the Mouse High Sensitive Fibroblast Growth Factor 15 ELISA Kit (FGF15) (ABclonal, RK02801) following the manufacturer’s protocols.

### Isolation of HSCs

As detailed in the literature [63], this process combines liver in situ retrograde pronase-collagenase perfusion with in vitro further digestion and density gradient-based separation of HSCs. Initially, the mouse was anesthetized via inhalation of isoflurane and cannulated at the inferior vena cava (IVC). Subsequently, the portal vein was cut and the liver was perfused sequentially with EGTA, pronase, and collagenase solutions. Following perfusion, the liver was excised and minced thoroughly in a petri dish. Further digestion was conducted in vitro using pronase/collagenase solution. The resulting liver cell suspension was filtered to remove undigested debris and washed with GBSS/B to remove excess digestion enzymes. The liver cells were then resuspended in density gradient medium (Nycodenz), and GBSS/B was slowly overlaid onto the cell-Nycodenz mixture. The gradient was centrifuged without braking, resulting in the formation of discontinuous layers. The HSC-containing layer at the top of the gradient was collected and centrifuged to obtain an HSC pellet, which was stored at −80°C for future use.

### Isolation of IECs

IECs were isolated from the entire small intestine (duodenum, jejunum, and ileum) by incubating the small intestine in 15 mM EDTA for 15 min. IECs were concentrated by centrifugation and the pellets were frozen [64].

### Serum ALT and AST measurement

The whole blood was clotted and centrifuged at 2,000 × g for 10 min at 4°C in a refrigerated benchtop microcentrifuge. The serum was transferred to a microcentrifuge tube for subsequent analysis. AST and ALT levels were measured using a colorimetric kit following established protocols at the BCM Mouse Metabolism and Phenotyping Core [65].

### Statistical analysis

Results are shown as the mean ± SEM. For experiments with only two groups, a Student’s *t*-test was used. For studies with three or more groups, one-way ANOVA was used, and for experiments with several groups with a balanced distribution of two factors, two-way ANOVA was used. A Sidak test was used for post-hoc analysis of comparisons within subgroups; *p* values < 0.05 were considered statistically significant. Additional details are provided in figure legends.

## Study approval

Animal care and experimental protocols were approved by the Institutional Animal Care and Use Committee of the Baylor College of Medicine.

## Data availability

All data supporting the findings of this study are available upon reasonable request. Mouse models are available from the corresponding author on request.

## Acknowledgments

We thank Mingyang Huang and Sarah Ishtiaq for their assistance with animal breeding, genotyping, and chemical assays.

## Author Contributions

Conceptualization, Y.Z.; investigation and analysis, X.C., H.L., and Y.D.; interpretation of data, X.C., S.Z., C.R.W., B.D., D.G., C.W., and Y.Z.; original draft preparation, X.C., J.M., and Y.Z.; reviewing and editing, all authors; supervision, Y.Z.; funding acquisition, Y.Z. All authors have read and agreed to the published version of the manuscript.

## Abbreviations

4-MU: 4-Methylumbelliferone
CDAHFD: choline-deficient, amino acid-defined, high-fat diet
MASH: metabolic dysfunction-associated steatohepatitis
BA: bile acid
FXR: farnesoid X receptor
HA: hyaluronan
GAG: glycosaminoglycan
GlcUA: D-glucuronic acid
GlcNAc: N-acetyl-D-glucosamine
MASH: nonalcoholic steatohepatitis
NAFLD: nonalcoholic fatty liver disease
MASH: metabolic dysfunction-associated steatohepatitis
FDA: Food and Drug Administration
THRβ: oral thyroid hormone receptor-beta
PIIINP: procollagen III N-terminal peptide
TIMP1: tissue inhibitor of metalloproteinase 1
ELF: enhanced liver fibrosis
HSCs: hepatic stellate cells
BDL: bile duct ligation
ECM: extracellular matrix
UGT: UDP-glycosyltransferases
IECs: intestinal epithelial cells
MCD: methionine/choline-deficient diet
PEG-PH20: pegylated human PH20
DIO: diet-induced obese
LPS: lipopolysaccharide
IVC: inferior vena cava

## References

[1] R.M. Hoffmann, G. Schwarz, C. Pohl, D.J. Ziegenhagen, W. Kruis, [Bile acid-independent effect of hymecromone on bile secretion and common bile duct motility], Dtsch Med Wochenschr 130(34-35) (2005) 1938–43.

[2] H.W. Krawzak, H.P. Heistermann, K. Andrejewski, G. Hohlbach, Postprandial bile-duct kinetics under the influence of 4-methylumbelliferone (hymecromone), Int J Clin Pharmacol Ther 33(10) (1995) 569–72.

[3] Y. Zhu, C. Crewe, P.E. Scherer, Hyaluronan in adipose tissue: Beyond dermal filler and therapeutic carrier, Sci Transl Med 8(323) (2016) 323ps4.

[4] Y. Zhu, I.L. Kruglikov, Y. Akgul, P.E. Scherer, Hyaluronan in adipogenesis, adipose tissue physiology and systemic metabolism, Matrix Biol (2018).

[5] I. Kakizaki, K. Kojima, K. Takagaki, M. Endo, R. Kannagi, M. Ito, Y. Maruo, H. Sato, T. Yasuda, S. Mita, K. Kimata, N. Itano, A novel mechanism for the inhibition of hyaluronan biosynthesis by 4-methylumbelliferone, J Biol Chem 279(32) (2004) 33281–9.

[6] A. Kultti, S. Pasonen-Seppanen, M. Jauhiainen, K.J. Rilla, R. Karna, E. Pyoria, R.H. Tammi, M.I. Tammi, 4-Methylumbelliferone inhibits hyaluronan synthesis by depletion of cellular UDP-glucuronic acid and downregulation of hyaluronan synthase 2 and 3, Exp Cell Res 315(11) (2009) 1914–23.

[7] S. Pouwels, N. Sakran, Y. Graham, A. Leal, T. Pintar, W. Yang, R. Kassir, R. Singhal, K. Mahawar, D. Ramnarain, Non-alcoholic fatty liver disease (NAFLD): a review of pathophysiology, clinical management and effects of weight loss, BMC Endocr Disord 22(1) (2022) 63.

[8] J.M. Fraile, S. Palliyil, C. Barelle, A.J. Porter, M. Kovaleva, Non-Alcoholic Steatohepatitis (NASH) - A Review of a Crowded Clinical Landscape, Driven by a Complex Disease, Drug Des Devel Ther 15 (2021) 3997–4009.

[9] A.C. Sheka, O. Adeyi, J. Thompson, B. Hameed, P.A. Crawford, S. Ikramuddin, Nonalcoholic Steatohepatitis: A Review, JAMA 323(12) (2020) 1175–1183.

[10] M.E. Rinella, J.V. Lazarus, V. Ratziu, S.M. Francque, A.J. Sanyal, F. Kanwal, D. Romero, M.F. Abdelmalek, Q.M. Anstee, J.P. Arab, M. Arrese, R. Bataller, U. Beuers, J. Boursier, E. Bugianesi, C.D. Byrne, G.E.C. Narro, A. Chowdhury, H. Cortez-Pinto, D.R. Cryer, K. Cusi, M. El-Kassas, S. Klein, W. Eskridge, J. Fan, S. Gawrieh, C.D. Guy, S.A. Harrison, S.U. Kim, B.G. Koot, M. Korenjak, K.V. Kowdley, F. Lacaille, R. Loomba, R. Mitchell-Thain, T.R. Morgan, E.E. Powell, M. Roden, M. Romero-Gomez, M. Silva, S.P. Singh, S.C. Sookoian, C.W. Spearman, D. Tiniakos, L. Valenti, M.B. Vos, V.W. Wong, S. Xanthakos, Y. Yilmaz, Z. Younossi, A. Hobbs, M. Villota-Rivas, P.N. Newsome, N.N.c. group, A multisociety Delphi consensus statement on new fatty liver disease nomenclature, Ann Hepatol 29(1) (2024) 101133.

[11] Z.M. Younossi, A.B. Koenig, D. Abdelatif, Y. Fazel, L. Henry, M. Wymer, Global epidemiology of nonalcoholic fatty liver disease-Meta-analytic assessment of prevalence, incidence, and outcomes, Hepatology 64(1) (2016) 73–84.

[12] S.L. Friedman, B.A. Neuschwander-Tetri, M. Rinella, A.J. Sanyal, Mechanisms of NAFLD development and therapeutic strategies, Nat Med 24(7) (2018) 908–922.

[13] L. Servin-Abad, E.R. Schiff, The Treatment of Hepatic Fibrosis: Reversal of the Underlying Disease Process, Gastroenterol Hepatol (N Y) 2(11) (2006) 819–825.

[14] K. Kingwell, NASH field celebrates ‘hurrah moment’ with a first FDA drug approval for the liver disease, Nat Rev Drug Discov (2024).

[15] R. Lichtinghagen, D. Pietsch, H. Bantel, M.P. Manns, K. Brand, M.J. Bahr, The Enhanced Liver Fibrosis (ELF) score: normal values, influence factors and proposed cut-off values, J Hepatol 59(2) (2013) 236–42.

[16] Y.M. Yang, M. Noureddin, C. Liu, K. Ohashi, S.Y. Kim, D. Ramnath, E.E. Powell, M.J. Sweet, Y.S. Roh, I.F. Hsin, N. Deng, Z. Liu, J. Liang, E. Mena, D. Shouhed, R.F. Schwabe, D. Jiang, S.C. Lu, P.W. Noble, E. Seki, Hyaluronan synthase 2-mediated hyaluronan production mediates Notch1 activation and liver fibrosis, Sci Transl Med 11(496) (2019).

[17] I.N. Andreichenko, A.A. Tsitrina, A.V. Fokin, A.I. Gabdulkhakova, D.I. Maltsev, G.S. Perelman, E.V. Bulgakova, A.M. Kulikov, A.S. Mikaelyan, Y.V. Kotelevtsev, 4-methylumbelliferone Prevents Liver Fibrosis by Affecting Hyaluronan Deposition, FSTL1 Expression and Cell Localization, Int J Mol Sci 20(24) (2019).

[18] Y.M. Yang, Z. Wang, M. Matsuda, E. Seki, Inhibition of hyaluronan synthesis by 4-methylumbelliferone ameliorates non-alcoholic steatohepatitis in choline-deficient L-amino acid-defined diet-induced murine model, Arch Pharm Res 44(2) (2021) 230–240.

[19] N. Sawada, K. Mikami, G. Igarashi, T. Endo, K. Sato, I. Kakizaki, S. Fukuda, Beneficial effect of 4-methylumbelliferone against bile duct ligation-induced hepatic fibrosis in rats, Hirosaki Medical Journal 66(2-4) (2016) 143–151.

[20] M. Kotulkar, D.R. Robarts, K. Lin-Rahardja, T. McQuillan, J. Surgnier, S.E. Tague, M. Czerwinski, K.L. Dennis, M.T. Pritchard, Hyaluronan synthesis inhibition normalizes ethanol-enhanced hepatic stellate cell activation, Alcohol Clin Exp Res (Hoboken) 47(8) (2023) 1544–1559.

[21] N. Halimani, M. Nesterchuk, A.A. Tsitrina, M. Sabirov, I.N. Andreichenko, N.O. Dashenkova, E. Petrova, A.M. Kulikov, T.S. Zatsepin, R.A. Romanov, A.S. Mikaelyan, Y.V. Kotelevtsev, Knockdown of Hyaluronan synthase 2 suppresses liver fibrosis in mice via induction of transcriptomic changes similar to 4MU treatment, Sci Rep 14(1) (2024) 2797.

[22] Y. Zhu, N. Li, M. Huang, M. Bartels, S. Dogne, S. Zhao, X. Chen, C. Crewe, L. Straub, L. Vishvanath, Z. Zhang, M. Shao, Y. Yang, C.M. Gliniak, R. Gordillo, G.I. Smith, W.L. Holland, R.K. Gupta, B. Dong, N. Caron, Y. Xu, Y. Akgul, S. Klein, P.E. Scherer, Adipose tissue hyaluronan production improves systemic glucose homeostasis and primes adipocytes for CL 316,243-stimulated lipolysis, Nat Commun 12(1) (2021) 4829.

[23] S. Wang, Q. Zhu, G. Liang, T. Franks, M. Boucher, K.K. Bence, M. Lu, C.M. Castorena, S. Zhao, J.K. Elmquist, P.E. Scherer, J.D. Horton, Cannabinoid receptor 1 signaling in hepatocytes and stellate cells does not contribute to NAFLD, J Clin Invest 131(22) (2021).

[24] S. Reitinger, G.T. Laschober, C. Fehrer, B. Greiderer, G. Lepperdinger, Mouse testicular hyaluronidase-like proteins SPAM1 and HYAL5 but not HYALP1 degrade hyaluronan, Biochem J 401(1) (2007) 79–85.

[25] L. Kang, L. Lantier, A. Kennedy, J.S. Bonner, W.H. Mayes, D.P. Bracy, L.H. Bookbinder, A.H. Hasty, C.B. Thompson, D.H. Wasserman, Hyaluronan accumulates with high-fat feeding and contributes to insulin resistance, Diabetes 62(6) (2013) 1888–96.

[26] X. Zhou, L. Cao, C. Jiang, Y. Xie, X. Cheng, K.W. Krausz, Y. Qi, L. Sun, Y.M. Shah, F.J. Gonzalez, G. Wang, H. Hao, PPARalpha-UGT axis activation represses intestinal FXR-FGF15 feedback signalling and exacerbates experimental colitis, Nat Commun 5 (2014) 4573.

[27] Y.C. Kim, M. Qi, X. Dong, S. Seok, H. Sun, B. Kemper, T. Fu, J.K. Kemper, Transgenic mice lacking FGF15/19-SHP phosphorylation display altered bile acids and gut bacteria, promoting nonalcoholic fatty liver disease, J Biol Chem 299(8) (2023) 104946.

[28] S. Ji, Q. Liu, S. Zhang, Q. Chen, C. Wang, W. Zhang, C. Xiao, Y. Li, C. Nian, J. Li, J. Li, J. Geng, L. Hong, C. Xie, Y. He, X. Chen, X. Li, Z.Y. Yin, H. You, K.H. Lin, Q. Wu, C. Yu, R.L. Johnson, L. Wang, L. Chen, F. Wang, D. Zhou, FGF15 Activates Hippo Signaling to Suppress Bile Acid Metabolism and Liver Tumorigenesis, Dev Cell 48(4) (2019) 460–474 e9.

[29] Y.C. Kim, S. Seok, Y. Zhang, J. Ma, B. Kong, G. Guo, B. Kemper, J.K. Kemper, Intestinal FGF15/19 physiologically repress hepatic lipogenesis in the late fed-state by activating SHP and DNMT3A, Nat Commun 11(1) (2020) 5969.

[30] S. Maliha, G.L. Guo, Farnesoid X receptor and fibroblast growth factor 15/19 as pharmacological targets, Liver Res 5(3) (2021) 142–150.

[31] A.-N. Moon, F. Briand, N. Breyner, D.-K. Song, M.R. Madsen, H. Kim, K. Choi, Y. Lee, W. Namkung, Improvement of NASH and liver fibrosis through modulation of the gut-liver axis by a novel intestinal FXR agonist, Biomedicine & Pharmacotherapy 173 (2024) 116331.

[32] C.J. Sinal, M. Tohkin, M. Miyata, J.M. Ward, G. Lambert, F.J. Gonzalez, Targeted disruption of the nuclear receptor FXR/BAR impairs bile acid and lipid homeostasis, Cell 102(6) (2000) 731–44.

[33] R. Meech, D.G. Hu, R.A. McKinnon, S.N. Mubarokah, A.Z. Haines, P.C. Nair, A. Rowland, P.I. Mackenzie, The UDP-Glycosyltransferase (UGT) Superfamily: New Members, New Functions, and Novel Paradigms, Physiol Rev 99(2) (2019) 1153–1222.

[34] C.L. Hsu, B. Schnabl, The gut-liver axis and gut microbiota in health and liver disease, Nat Rev Microbiol 21(11) (2023) 719–733.

[35] N.Y. Lee, K.T. Suk, The Role of the Gut Microbiome in Liver Cirrhosis Treatment, Int J Mol Sci 22(1) (2020).

[36] X. Ge, C. Ding, W. Zhao, L. Xu, H. Tian, J. Gong, M. Zhu, J. Li, N. Li, Antibiotics-induced depletion of mice microbiota induces changes in host serotonin biosynthesis and intestinal motility, J Transl Med 15(1) (2017) 13.

[37] G. Rizzatti, L.R. Lopetuso, G. Gibiino, C. Binda, A. Gasbarrini, Proteobacteria: A Common Factor in Human Diseases, Biomed Res Int 2017 (2017) 9351507.

[38] E. Biagi, L. Nylund, M. Candela, R. Ostan, L. Bucci, E. Pini, J. Nikkila, D. Monti, R. Satokari, C. Franceschi, P. Brigidi, W. De Vos, Through ageing, and beyond: gut microbiota and inflammatory status in seniors and centenarians, PLoS One 5(5) (2010) e10667.

[39] T. Sen, C.R. Cawthon, B.T. Ihde, A. Hajnal, P.M. DiLorenzo, C.B. de La Serre, K. Czaja, Diet-driven microbiota dysbiosis is associated with vagal remodeling and obesity, Physiol Behav 173 (2017) 305–317.

[40] J. Qin, Y. Li, Z. Cai, S. Li, J. Zhu, F. Zhang, S. Liang, W. Zhang, Y. Guan, D. Shen, Y. Peng, D. Zhang, Z. Jie, W. Wu, Y. Qin, W. Xue, J. Li, L. Han, D. Lu, P. Wu, Y. Dai, X. Sun, Z. Li, A. Tang, S. Zhong, X. Li, W. Chen, R. Xu, M. Wang, Q. Feng, M. Gong, J. Yu, Y. Zhang, M. Zhang, T. Hansen, G. Sanchez, J. Raes, G. Falony, S. Okuda, M. Almeida, E. LeChatelier, P. Renault, N. Pons, J.M. Batto, Z. Zhang, H. Chen, R. Yang, W. Zheng, S. Li, H. Yang, J. Wang, S.D. Ehrlich, R. Nielsen, O. Pedersen, K. Kristiansen, J. Wang, A metagenome-wide association study of gut microbiota in type 2 diabetes, Nature 490(7418) (2012) 55–60.

[41] J.T. Haas, S. Francque, B. Staels, Pathophysiology and Mechanisms of Nonalcoholic Fatty Liver Disease, Annu Rev Physiol 78 (2016) 181–205.

[42] R. Loomba, S.L. Friedman, G.I. Shulman, Mechanisms and disease consequences of nonalcoholic fatty liver disease, Cell 184(10) (2021) 2537–2564.

[43] S. Tang, J. Borlak, Genomics of human NAFLD: Lack of data reproducibility and high interpatient variability in drug target expression as major causes of drug failures, Hepatology 80(4) (2024) 901–915.

[44] S.A. Harrison, A.M. Allen, J. Dubourg, M. Noureddin, N. Alkhouri, Challenges and opportunities in NASH drug development, Nat Med 29(3) (2023) 562–573.

[45] B.E. Harvey, NASH: regulatory considerations for clinical drug development and U.S. FDA approval, Acta Pharmacol Sin 43(5) (2022) 1210–1214.

[46] E. Guirguis, J. Dougherty, K. Thornby, Y. Grace, K. Mack, Resmetirom: The First Food and Drug Administration-Approved Medication for Nonalcoholic Steatohepatitis (NASH), Ann Pharmacother 59(2) (2025) 162–173.

[47] S. Reagan-Shaw, M. Nihal, N. Ahmad, Dose translation from animal to human studies revisited, FASEB J 22(3) (2008) 659–61.

[48] A. Rowland, J.O. Miners, P.I. Mackenzie, The UDP-glucuronosyltransferases: their role in drug metabolism and detoxification, Int J Biochem Cell Biol 45(6) (2013) 1121–32.

[49] J.I. Rosser, N. Nagy, R. Goel, G. Kaber, S. Demirdjian, J. Saxena, J.B. Bollyky, A.R. Frymoyer, A.E. Pacheco-Navarro, E.B. Burgener, J. Rajadas, Z. Wang, O. Arbach, C.E. Dunn, A. Kalinowski, C.E. Milla, P.L. Bollyky, Oral hymecromone decreases hyaluronan in human study participants, J Clin Invest 132(9) (2022).

[50] B. McDonald, E.F. McAvoy, F. Lam, V. Gill, C. de la Motte, R.C. Savani, P. Kubes, Interaction of CD44 and hyaluronan is the dominant mechanism for neutrophil sequestration in inflamed liver sinusoids, J Exp Med 205(4) (2008) 915–27.

[51] B.P. Toole, Hyaluronan: from extracellular glue to pericellular cue, Nat Rev Cancer 4(7) (2004) 528–39.

[52] M.G. Neuman, R.M. Nanau, L. Oruna-Sanchez, G. Coto, Hyaluronic acid and wound healing, J Pharm Pharm Sci 18(1) (2015) 53–60.

[53] G.J. Mulder, S. Brouwer, J.G. Weitering, E. Scholtens, K.S. Pang, Glucuronidation and sulfation in the rat in vivo. The role of the liver and the intestine in the in vivo clearance of 4-methylumbelliferone, Biochem Pharmacol 34(8) (1985) 1325–9.

[54] B. Madhoolika, N.V. Anil Kumar, S. Balaji, In vitro analysis of 4-methylumbelliferone as a sole carbon source for Lactobacillus helveticus 2126, Lett Appl Microbiol 65(3) (2017) 249–255.

[55] G. Marroncini, L. Naldi, S. Martinelli, A. Amedei, Gut-Liver-Pancreas Axis Crosstalk in Health and Disease: From the Role of Microbial Metabolites to Innovative Microbiota Manipulating Strategies, Biomedicines 12(7) (2024).

[56] K. Sun, I. Wernstedt Asterholm, C.M. Kusminski, A.C. Bueno, Z.V. Wang, J.W. Pollard, R.A. Brekken, P.E. Scherer, Dichotomous effects of VEGF-A on adipose tissue dysfunction, Proc Natl Acad Sci U S A 109(15) (2012) 5874–9.

[57] C. Postic, M. Shiota, K.D. Niswender, T.L. Jetton, Y. Chen, J.M. Moates, K.D. Shelton, J. Lindner, A.D. Cherrington, M.A. Magnuson, Dual roles for glucokinase in glucose homeostasis as determined by liver and pancreatic beta cell-specific gene knock-outs using Cre recombinase, J Biol Chem 274(1) (1999) 305–15.

[58] G. Belteki, J. Haigh, N. Kabacs, K. Haigh, K. Sison, F. Costantini, J. Whitsett, S.E. Quaggin, A. Nagy, Conditional and inducible transgene expression in mice through the combinatorial use of Cre-mediated recombination and tetracycline induction, Nucleic Acids Res 33(5) (2005) e51.

[59] F. el Marjou, K.P. Janssen, B.H. Chang, M. Li, V. Hindie, L. Chan, D. Louvard, P. Chambon, D. Metzger, S. Robine, Tissue-specific and inducible Cre-mediated recombination in the gut epithelium, Genesis 39(3) (2004) 186–93.

[60] L. Xu, A.Y. Hui, E. Albanis, M.J. Arthur, S.M. O’Byrne, W.S. Blaner, P. Mukherjee, S.L. Friedman, F.J. Eng, Human hepatic stellate cell lines, LX-1 and LX-2: new tools for analysis of hepatic fibrosis, Gut 54(1) (2005) 142–51.

[61] Y. Zhu, K.M. Pires, K.J. Whitehead, C.D. Olsen, B. Wayment, Y.C. Zhang, H. Bugger, O. Ilkun, S.E. Litwin, G. Thomas, S.C. Kozma, E.D. Abel, Mechanistic target of rapamycin (Mtor) is essential for murine embryonic heart development and growth, PLoS One 8(1) (2013) e54221.

[62] X. Wang, A. Spandidos, H. Wang, B. Seed, PrimerBank: a PCR primer database for quantitative gene expression analysis, 2012 update, Nucleic Acids Res 40(Database issue) (2012) D1144–9.

[63] I. Mederacke, D.H. Dapito, S. Affo, H. Uchinami, R.F. Schwabe, High-yield and high-purity isolation of hepatic stellate cells from normal and fibrotic mouse livers, Nat Protoc 10(2) (2015) 305–15.

[64] M. Davalos-Salas, M.K. Montgomery, C.M. Reehorst, R. Nightingale, I. Ng, H. Anderton, S. Al-Obaidi, A. Lesmana, C.M. Scott, P. Ioannidis, H. Kalra, S. Keerthikumar, L. Togel, A. Rigopoulos, S.J. Gong, D.S. Williams, P. Yoganantharaja, K. Bell-Anderson, S. Mathivanan, Y. Gibert, S. Hiebert, A.M. Scott, M.J. Watt, J.M. Mariadason, Deletion of intestinal Hdac3 remodels the lipidome of enterocytes and protects mice from diet-induced obesity, Nat Commun 10(1) (2019) 5291.

[65] X. Chen, H. Li, Y. Liu, J. Qi, B. Dong, S. Huang, S. Zhao, Y. Zhu, Dimethyl Sulfoxide Inhibits Bile Acid Synthesis in Healthy Mice but Does Not Protect Mice from Bile-Acid-Induced Liver Damage, Biology (Basel) 12(8) (2023).

