## Supplementary material for "The potential of 4-Methylumbelliferone to be repurposed for treating liver fibrosis"

### Mechanisms by which 4-Methylumbelliferone inhibits liver fibrogenesis

Table of Contents

Supplementary figures	2

### Supplementary Figures S1

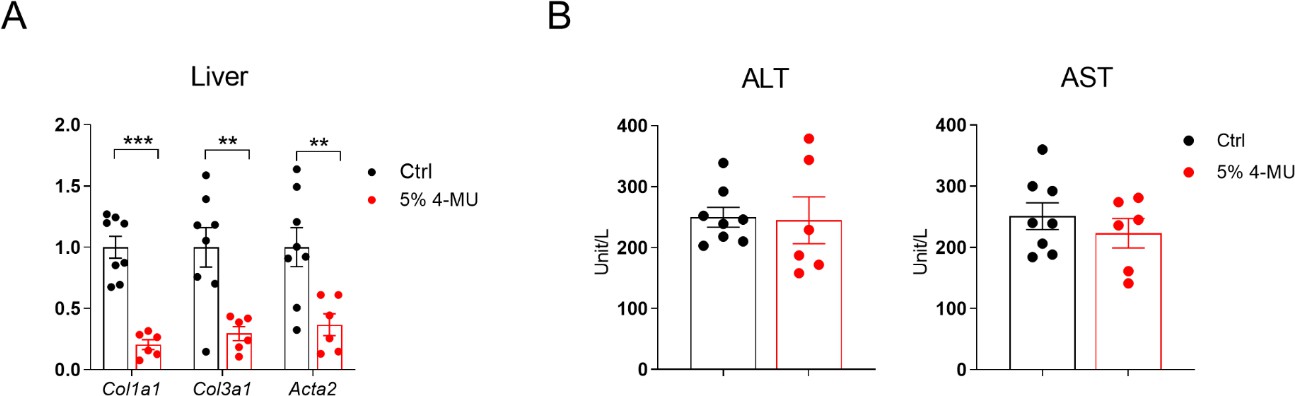

**Supplementary Fig. 1. (A)** Relative mRNA expression of fibrosis-related genes in the livers of C57BL/6J mice kept on CDAHFD or CDAHFD with 5% 4-MU for 8 weeks. **(B)** Serum ALT and AST levels of C57BL/6J mice from panel A; for panels A-B, n = 6-8. All data are presented as mean ± SEM; * *p* < 0.05, ** *p* < 0.01, *** *p* < 0.001.

# S2

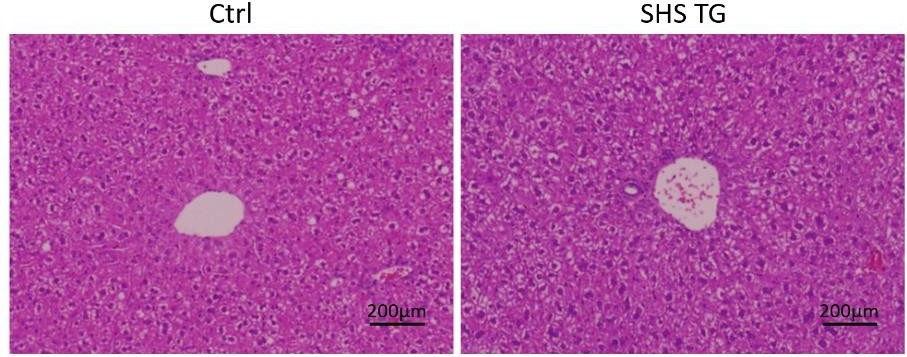

**Supplementary Fig. 2.** Representative images of liver H&E staining in control and SHS TG mice fed with 4-week dox200 MCD diet, scale bar = 200 μm.

# S3

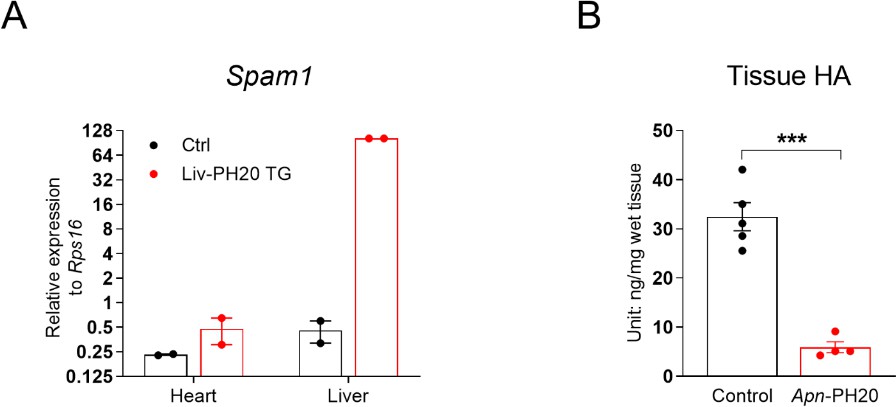

**Supplementary Fig. 3. (A)** mRNA expression of *Spam1* in heart and liver lysates from Alb-cre::Rosa-rtTA (Ctrl) or Alb-Cre:Rosa-rtTA::TRE-PH20 (Liv-PH20 TG) mice treated with normal chow (NC) containing 200 mg/kg doxycycline for 5 days, n = 2. **(B)** iWAT HA content of *Apn*-rtTA (control) or *Apn*-rtTA::TRE-PH20 (referred to as *Apn*-PH20) mice fed with dox200 NC for 5 days, n = 4-5. All data are presented as mean ± SEM; *** *p* < 0.001.

# S4

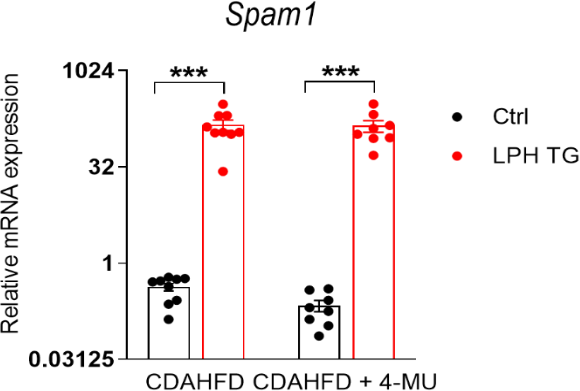

**Supplementary Fig. 4.** mRNA expression of *Spam1* gene in the livers of control and SPH TG mice kept on CDAHFD with or without 5% 4-MU for 8 weeks. All data are presented as mean ± SEM; *** *p* < 0.001.

# S5

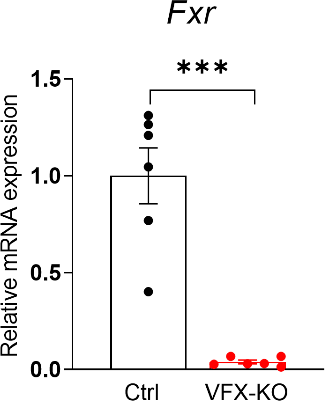

**Supplementary Fig. 5.** Relative mRNA expression of *Fxr* in isolated IECs from control and VFX-KO mice after one injection of tamoxifen (100 mg/kg body weight, intraperitoneal injection), n = 6. All data are presented as mean ± SEM; *** *p* < 0.001.

# S6

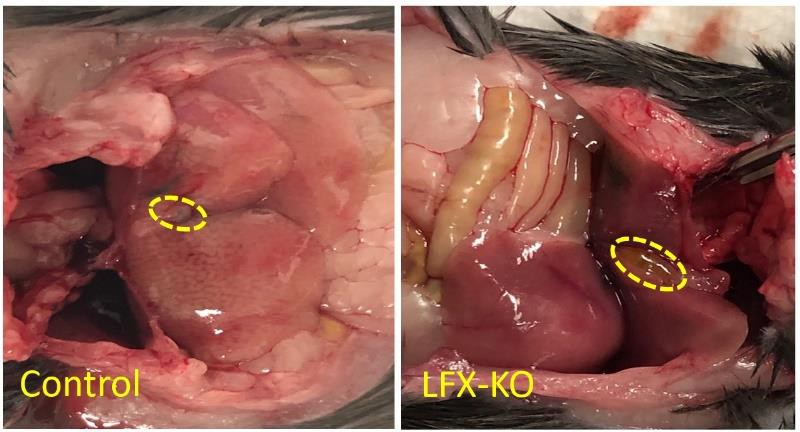

**Supplementary Fig 6** Abdominal gross appearance of control and LFX-KO mice fed with 8-week HFD.

# S7

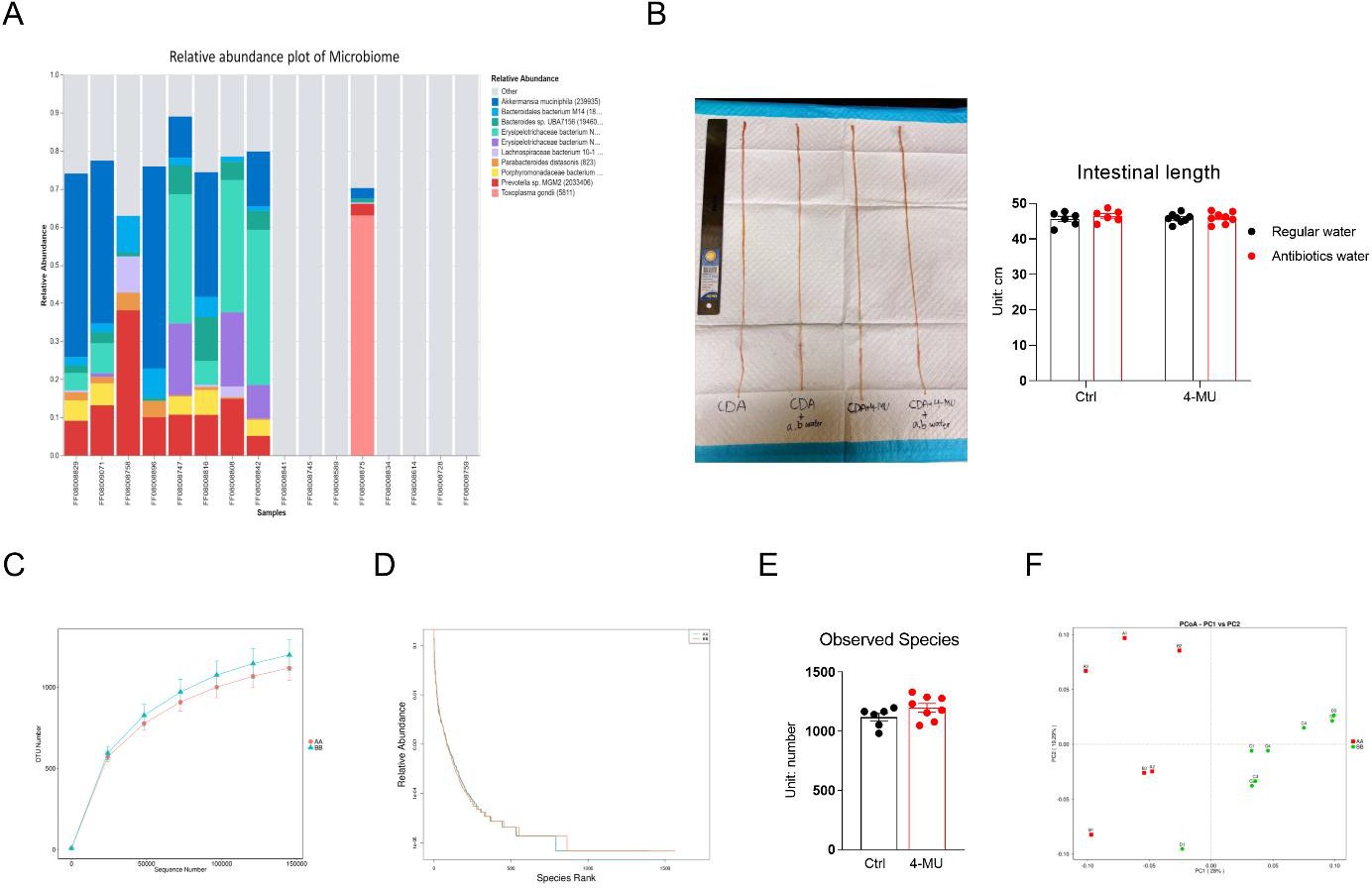

**Supplementary Fig. 7. (A)** Relative abundance plot of microbiome in WT mice treated with antibiotic water for 3 weeks (eight samples on the right); regular water was used as control (eight samples on the left). There was Toxoplasma gondii (parasite) infection in one antibiotic water-treated mouse. **(B)** Left: Gross appearance of whole intestine of mice referred to in Fig. 7A. Right: Whole intestine length of mice referred to in Fig. 7A, n = 6-8. **(C)** Species accumulation curves; and **(D)** rank abundance curves of gut microbiota community structures of AA (CDAHFD) and BB (4-MU + CDAHFD) groups. **(E)** Alpha diversity expressed as observed species in Ctrl and 4-MU groups, n = 6-8. **(F)** Principal coordinate analysis (PCoA) based on weighted Unifrac distance. All data are presented as mean ± SEM.
